# The electric field generated by cationic TiO_2_ nanoparticles can be their key antimicrobial factor

**DOI:** 10.1101/2022.05.23.492822

**Authors:** Lyudmila V. Zhukova

## Abstract

The antimicrobial effect of TiO_2_ nanoparticles (TiO_2_ NPs) has been reliably established. However, the mechanism of this action remains unclear. Currently, according to the dominant hypothesis, it is assumed that the key destructive mechanism is the photocatalytic oxidation of cell components, and the key antimicrobial factor is the reactive oxygen species (ROS) formed on the surface of TiO_2_ NPs as a result of photocatalysis. In this communication, for the first time, the influence of the electric field created by cationic TiO_2_ NPs is suggested as a key killing factor, and the key antimicrobial mechanism - the electrical breakdown of the cell membranes and the destruction of the DNA molecule under the action of this field. Here, it was shown by mathematical calculation that TiO_2_ NPs are able to generate an electric field with a strength sufficient for an electrical breakdown of bilayer lipid membranes, which can lead to a violation of the structure and vital functions of the cytoplasmic membrane (CPM) and to collapse of DNA molecule.

**GRAPHICAL ABSTRACT:** 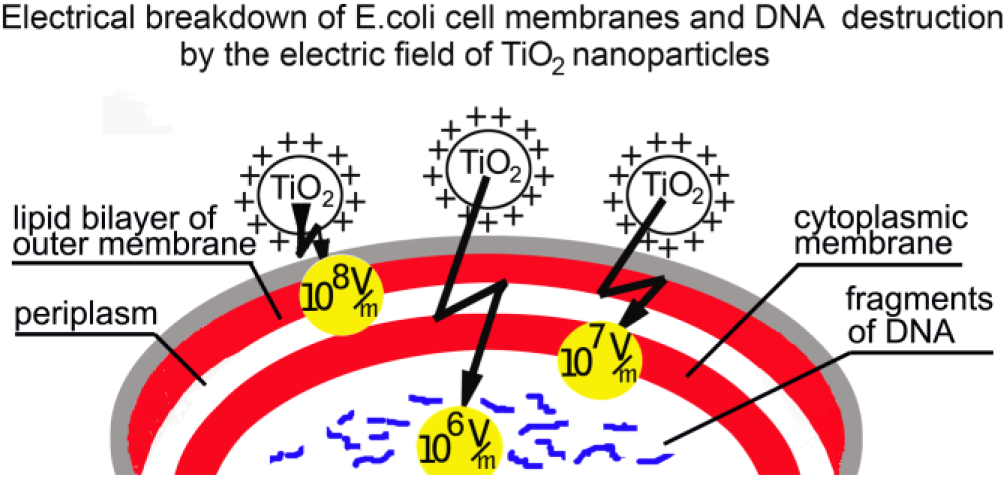

## 1. Introduction

At present, nanoparticles of inorganic semiconductor TiO_2_ NPs attract the attention of researchers as an alternative to antibiotics^1–3^. Facts have been obtained indicating that TiO_2_ NPs can have a destructive effect on microorganisms of various taxonomic groups^4–6^.

Initially, it was believed that only nanoparticles that absorbed radiation with an energy greater than or equal to the band gap energy could have such an effect^7,8^. For TiO_2_ NPs, this energy is approximately 3.2 eV. It means that it is enough to irradiate the TiO_2_ NPs with radiation of below 385 nm, i.e. near-ultra violet light (UV-A). However, later it was shown that even without UV-A irradiation, TiO_2_ NPs can have an antimicrobial effect^9–11^.

TiO_2_ NPs tend to change the surface charge in magnitude and sign depending on the pH value of the medium. Figure 1 shows the dependence of the zeta potential on the pH of the medium for TiO_2_ NPs of several commercial preparations, with the use of which a significant decrease in the number of colony-forming units (CFUs) was observed.

**Figure 1.**
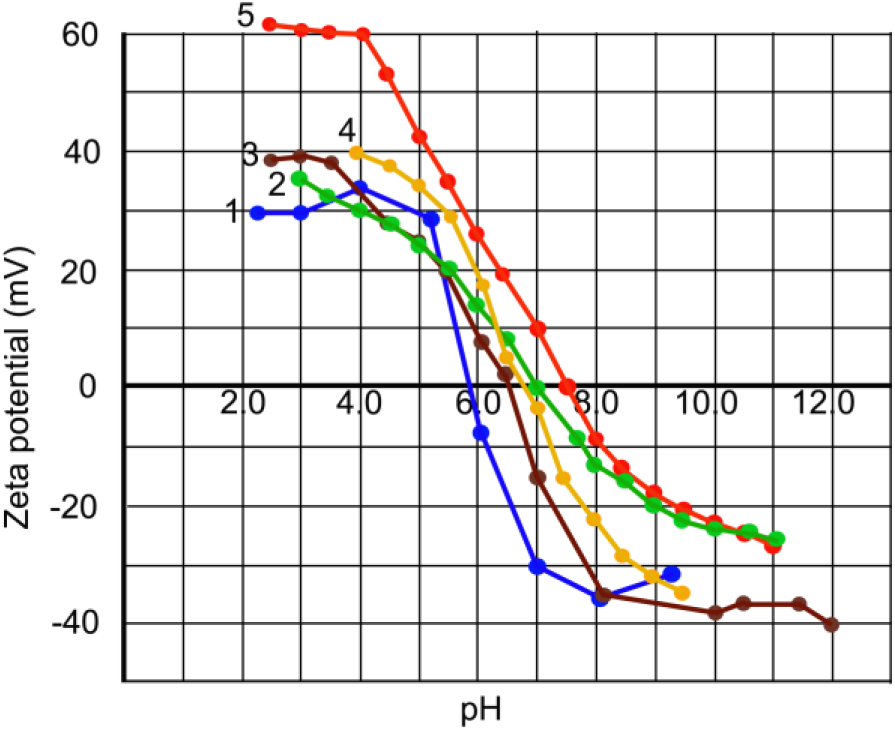
Dependence of the zeta potential of TiO_2_ NPs on the pH of the medium for commercial preparations: 1 - Degussa P 25^12^; 2 - Degussa P 25^13^; 3 – P 25 (Evonic Industries)^14^; 4 - Degussa P 25 TN90^13^; 5 - TKS 201 (Tayca Chemical, Japan)^10^.

Zeta-potential analysis showed that the isoelectric point (pI) of TiO_2_ suspensions (when they are not charged) in different preparations of these nanoparticles is near pH 6-7.5 and their surface is negatively charged at pH > 6.0 - 7.5 and positively charged at pH < 7.5 - 6.0. ^10,12–14^

Under pH conditions close to pI, when TiO_2_ NPs are practically uncharged, they spontaneously aggregate into larger clusters, while under acidic or alkaline conditions, a colloidal suspension of nanoparticles is more stable due to the repulsion of similarly charged nanoparticles.^12^

When cells were treated with TiO_2_ NPs in combination with UV-A irradiation, cell inactivation was observed both in the case when the nanoparticles were not charged, and in the case of their positive or negative charge.^13, 14^

When using TiO_2_ NPs in experiments without UV-A irradiation, the antimicrobial effect was observed under pH conditions below the pI when the TiO_2_ NPs were positively charged.^10,11^ At pH above pI, when TiO_2_ NPs are negatively charged, the action of TiO_2_ NPs did not lead to cell death.^11^ If the experiments were carried out in a neutral environment, that is, close to pI, when the TiO_2_ NPs were practically uncharged, the decrease in the number of the CFUs was not observed or was insignificant.^8,11,13,15^

With the above information, the mechanism of bacterial cell inactivation is still not understood fully. The main mechanisms of the cell inactivation described in the literature are photocatalytic oxidation^2,4–6,16^ and mechanisms not associated with photocatalysis, listed in publications.^1–3, 11^

Currently, photocatalytic oxidation is considered by most researchers as a key mechanism leading to the death of microorganisms. Non-oxidative mechanisms are given secondary importance. The hypothesis of the primary role of photocatalytic oxidation is based on the ability of these nanoparticles under the UV-A irradiation to form on their surface ROS, which is believed to damage cell components, starting from the cell wall and then, gradually going deeper into the cell, they destroy its contents, including the CPM and the DNA, which leads to the death of the microorganism.^2,4–6,15,16^

In the absence of UV-A irradiation, when ROS are not formed, it is assumed that cell death occurs due to direct contact of nanoparticles with the cell surface, which leads to disruption of the properties and functions of the cell membrane, in particular, its integrity, permeability, osmoregulation, interruption of electron transfer, a change in the transmembrane potential.^11,17,18^

It should be noted that when cells are exposed to TiO2 NPs without UV-A irradiation, one can obtain almost the same decrease in CFUs value (by several orders of magnitude)^10^ as when exposed in combination with UV-A irradiation^13^. This fact indicates that the effect of ROS formed during photocatalysis is not the key one, since nanoparticles can have almost the same detrimental effect even in the absence of ROS. This suggests that it is not ROS generated during photocatalysis that play a key role in the antimicrobial action, but some other factor, since in the absence of UV-A irradiation, photocatalysis does not occur and there is no formation of ROS associated with it.

In this article, a hypothesis has been put forward that suggests that the key killing factor is the influence of the electric field created by cationic TiO2 NPs, both in combination with UV-A irradiation or without it, which causes a violation of the structure and semipermeable properties of the CPM, collapse DNA, which ultimately leads to cell death. The author made this assumption on the basis of a comparative analysis of literature data on the effect of TiO2 NPs and an external electric field on bacterial cells, which shows that the response of cells to the action of TiO2 NPs is very similar to their response to the influence of an external electric field.

It was previously revealed that exposure to strong electric fields in the form of short pulses can have a lethal effect on the suspensions of microorganisms belonging to different taxonomic groups.^19,20^ The short pulses with relatively long intervals were used to minimize the temperature rise, in demonstrating the effect of the electric field itself.

It should be noted that a bacterial cell against the influence of an external electric field has powerful protection in the form of bilayer lipid membranes with powerful insulating properties. They have high electrical resistance and the ability to withstand very high electric fields. It is possible to overcome the electrical resistance of the membrane, that is, to cause a sharp increase in its conductivity if exposed to an electric field of several kilovolts per centimeter in the form of pulses lasting from tens of microseconds.^21^ Under the action of an external electric field the induction of the transmembrane potential occurs.^22^ The growth of the latter can reach a critical value, called the electrical breakdown potential, which in biological membranes of different objects is approximately equal to 1V. In particular, for E. coli spheroplasts the electrical breakdown potential is 0.85 V.^22^

When the transmembrane voltage induced by an external electric field reaches the electrical breakdown potential, the membrane loses the intrinsic properties, such as electrical resistance, membrane potential, barrier function, ets. A significant increase in the permeability of the cell membrane for ions, molecules, and even macromolecules occurs.^23^ These changes are explained by the formation and growth of pores in the membrane. Chang and Reese^24^, using fast freezing electron microscopy, have shown that membrane pores are indeed formed as a result of exposure to an electric field.

This phenomenon is based on the spontaneous generation of defects in the lipid bilayer due to the thermal motion of phospholipid molecules. In the absence of a potential difference across the membrane, an increase in the size of spontaneously formed pores does not occur. Under the action of surface tension forces, a spontaneously formed defect (pore) is immediately tightened and the membrane remains intact. With an increase in the potential difference across the membrane, the energy required for the formation and growth of a pore decreases. At a critical potential difference (electrical breakdown potential), the growth of formed pores becomes spontaneous.^25^

The short-term effect of an electric field on a bilayer lipid membrane is reversible. However, at a constant transmembrane potential, if it exceeds the critical value, the current spontaneously increases in time until the complete destruction of the membrane.^23,25^ It was revealed that a transmembrane potential close to 1 V (electrical breakdown potential) can generate an external electric field from several units to several tens of kV/cm,^21,22^ that is, of the order of 10^5^ V/m - 10^6^ V/m.

It should be noted that the outer membrane, which is part of the cell wall, is not a vital structure, since even complete removal of the cell wall does not lead to cell death. A cell with a completely removed cell wall (protoplast) retains the ability to carry out metabolism, including respiration, and the ability to grow.^26^ However, it is obvious that the outer membrane is an insulating barrier that protects the CPM from the effects of an external electric field.

Unlike the outer membrane, the CPM is a vital structure. The CPM in bacterial cells is energy-transforming.^27^ The respiratory chain of aerobic bacteria is located in the CPM. The transfer of electrons in the respiratory chain of the prokaryotes creates a difference in the electrochemical potential of H^+^ ions (ΔμH^+^), which includes two components - the transmembrane difference in the concentration of hydrogen ions (ΔpH) and the transmembrane difference in the electric potential (△Ψ). The presence of a transmembrane potential difference is a necessary condition for obtaining energy.

Energy in the form of ΔμH^+^ or its components can be used in various energy-dependent processes localized on the membrane: for chemical work (synthesis of ATP), for mechanical work (rotation of the flagellum), for osmotic work, which in a broad sense includes the trans-membrane transfer of matters to its higher concentration side and other functions.^27^ Besides, the CPM performs a barrier function. It is characterized by a pronounced selective permeability: with the help of special carriers, a selective transfer of various organic and inorganic molecules and ions is carried out through the membrane. It is the low permeability of the CPM in combination with the selective transport of ions and molecules that is a prerequisite for maintaining the transmembrane potential and non-equilibrium concentration of ions and molecules in the cytoplasm with respect to the external environment, which is a necessary condition for the course of life processes. Violation of the membrane structure, resulting in an increase in its permeability, leads to a decrease and, ultimately, to zeroing of the trans-membrane potential, as well as to an equalization of the concentration of ions and molecules inside and outside the cell, which means death for the cell. It should be noted that there are repeatedly confirmed facts indicating a violation of the semipermeable properties of CPM as a result of exposure of bacterial cells to TiO_2_ NPs in combination with or without UV-A irradiation^4,2,8,17,18,28^.

It should be expected that the destruction of bilayer lipid membranes should lead to the disappearance of the insulating barrier protecting the DNA from an external electric field. It is known that an electric field with a voltage of several tens of kilovolts per meter affects the structure and integrity of the DNA molecule. Tang et al.^29^ reported that an electric field of ~ 200 V/cm can induce a strong compression, spontaneous self-entanglement, and tying of polymer chains inside individual DNA molecules. Recent studies have revealed that in the presence of a strong electric field, a double-stranded DNA molecule can be destroyed. Using the comet assay method it has been demonstrated that an electric field up to 200 kV/m induces DNA fragmentation.^30^

According to Zhou et al.^31^, the critical voltage of the electric field at which DNA molecules collapse is of the order of 10^5^ V/m. The driving force of the collapse process is still not fully understood.^31^

If, indeed, when exposed to TiO_2_ NPs, cell death occurs as a result of the influence of the nanoparticles’ electric field, this means that the strength of this field should be sufficient to generate the electrical breakdown potential of the bilayer lipid membrane of bacterial cells, that is, on the order of 10^5^ - 10^6^ V/m.

The purpose of this communication is to put forward a hypothesis explaining the antimicrobial effect of TiO_2_ NPs by the influence of the electric field generated by these nanoparticles, as well as to clarify the question of whether the TiO_2_ NPs can generate an electric field with a strength sufficient to destroy the cell membrane and DNA molecule.

It is quite difficult to make an accurate calculation of the electric field strength generated by TiO_2_ NPs, since the nanoparticles covering the cell have different sizes and are distributed unevenly, which follows from their SEM images.^14^ Here, for an approximate estimation, a calculation of the electric field strength inside the cell, which can be created by a cationic TiO_2_ nanoparticle located on the surface of a bacterial cell at a distances, corresponding to the location of the outer membrane lipid bilayer, the CPM, as well as in the central part of the cell, where the DNA molecule is located was carried out.

## 2. Materials and methods

### 2. 1. Antimicrobial agent – TiO_2_ NPs

Calculations were carried out on the example of a nanoparticle of a commercial preparation P-25 (Degussa Co., Germany), often used in experiments to study the antimicrobial action of nanoparticles. The crystal structure and particulate properties for the P-25 TiO_2_ powders were studied by X-ray diffraction and transmission electron microscopy were described in publications.^11,12^ The phase structure of the P-25 powders is predominantly anatase (80% - 70%) mixed with a small amount of rutile (20% - 30%). The P-25 TiO_2_ particles are mono-dispersed and spherical with a diameter of 20-50 nm.^12,13^

According to different authors, the pI of P-25 TiO_2_ suspensions is near pH 6 – 7 (Figure 1). The zeta potential of P-25 TiO_2_ NPs sharply increases from 0 at pH 6 to about +30 mV at pH 5 remaining almost unchanged around (+30) – (+40) mV at lower pH values (down to 2) and drops sharply above pH 7 up to about −30 mV.

The hydrodynamic diameter of TiO_2_ particles is maximum at the pI. However, the suspension is a stable colloid at both acid and basic pH far from its pI point.^11,12^

For calculating the charge of TiO_2_ nanoparticles we choose the simplest case when the TiO_2_ nanoparticle has a spherical shape. In this case, the surface charge density is the same at all points on its surface. The surface charge density of TiO_2_ NPs depends on the pH value.^32^ We choose a pH value of 4.0, at which, as previously shown in our work,^10^ the number of CFUs of the E. coli K 12 cells decreases by four orders of magnitude during 60 minutes of dark exposure to TiO_2_ NPs. Under these conditions, the positive zeta potential of TiO_2_ NPs is close to the maximum (Figure 1) and the aggregation of nanoparticles is minimal due to their high surface charge.

Besides, at this pH value, the close contact between the surfaces of cells and nanoparticles takes place due to electrostatic interaction, because under this acidic condition the surface of TiO_2_ NPs is positively charged, and the bacterial cell surface is - negatively charged. As established by Holmberg et al.^32^, the surface charge density (σ) for TiO_2_ NPs of the commercial preparation P-25 at a pH of about 4.0 is approximately equal to 0.1 C/m^2^. Pagnout et al.^11^ found that the hydrodynamic diameter of TiO_2_ NPs at pH about 4.0 is approximately 80 nm.

### 2. 2. The outer cell membrane and the CPM of the E. coli K 12 cell- the object of the TiO_2_ NPs action

The gram-negative bacterium Escherichia coli K 12, which often is used as a test microorganism, was chosen for the calculation of electric field action.

Figure 2 shows a schematic image of the outer shell structure of the gram-negative bacterium, based on the data presented by Silhavy et al.^33^ and Matias et al.^34^ This figure gives us an idea of the cell wall components and the distance to the outer membrane lipid bilayer and the CPM from the cell surface.

**Figure 2.**
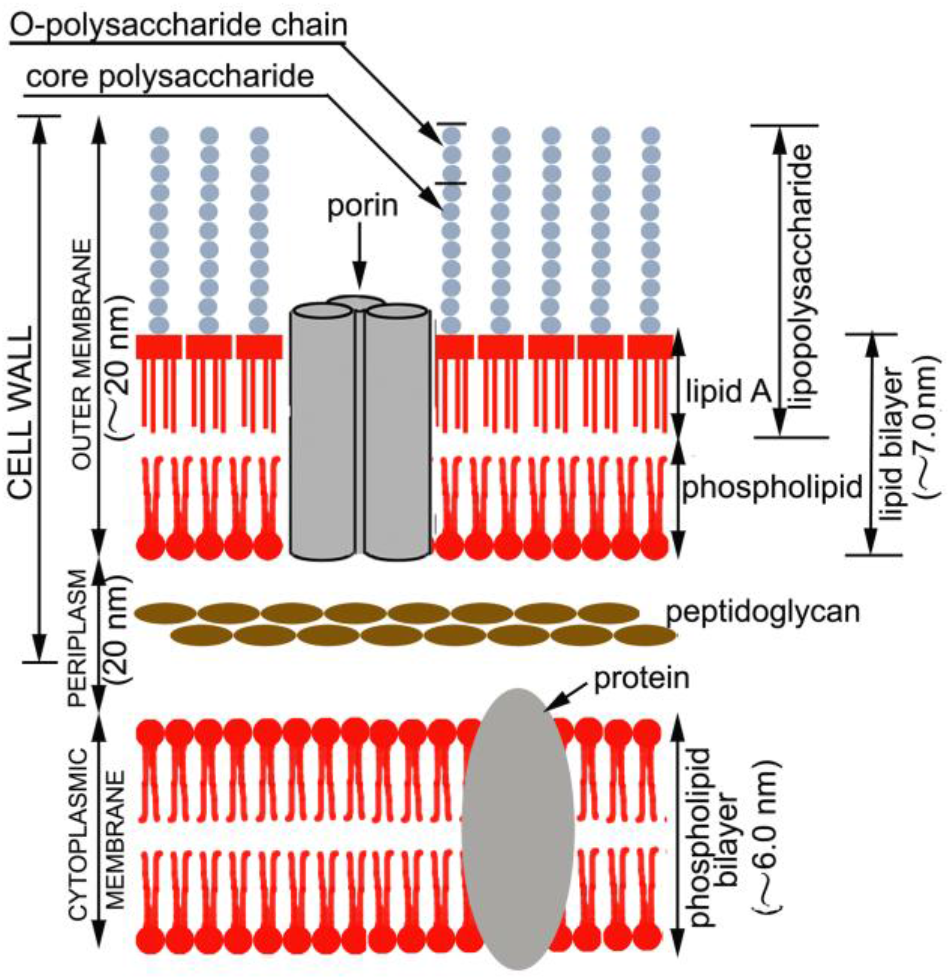
Schematic depiction of the outer shell of the Gram-negative bacterium.

In E. coli K 12, the outer membrane lipid bilayer is located relatively close to the cell surface, since there are no O-polysaccharide chains in this strain (Figure 1). As follows from the literature data, the thickness of the polysaccharide layer is no more than 10 nm, and the distance to the CPM is approximately 40 nm.

### 2. 3. Methods

The electric field strength E was calculated using the Coulomb formula derived for a point charge:

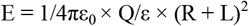

Where Q - the charge of the nanoparticle; R - the radius of the nanoparticle; L - the distance at which the electric field strength is determined; ε_0_ - electric constant equal to 8.85 × 10^-12^ (CN^-1^m^-2^); ε - dielectric constant of the medium equal to 80 (dimensionless value).

The surface area of the nanoparticle was calculated using the Archimedes formula derived to determine the surface area of a sphere:

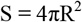

where S - the surface area of the nanoparticle; R - the radius of the nanoparticle; π - a mathematical constant equal to 3.14.

The charge of the nanoparticle was calculated by the equation:

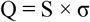

where Q - the charge of the nanoparticle; S - the surface area of the nanoparticle; σ - the surface charge density.

## 3. Results and discussion

In order to clarify the question of whether the electric field of TiO_2_ NPs can cause a violation of the integrity of the outer membrane lipid bilayer and the CPM of E. coli K 12 cells, it is possible to calculate the electric field strength that is created by the cationic TiO_2_ nanoparticle inside the cell at a distance of 10 nm and 40 nm, which is approximately equal to the distance from the surface to the outer membrane lipid bilayer and CPM correspondently (Figure 2). It is also possible to calculate the electric field strength at a distance of 460 nm, which is approximately equal to the distance from the lateral surface to the middle of the cell and the DNA location. Table 1 shows the results of the calculated electric field strength.

**TABLE 1.** Electric field strength E (V/m) inside the cell at different distances from the nanoparticle.

| The distance from the nanoparticle $L$ (nm), at which the electric field strength is determined | Electric field strength $E$ (V/m) |
| --- | --- |
| 10 | $9,0 \times 10^7$ ($\sim 10^8$) |
| 40 | $3,5 \times 10^7$ ($\sim 10^7$) |
| 460 | $9,0 \times 10^5$ ($\sim 10^6$) |

Thus, the electric field strength that a cationic TiO_2_ nanoparticle can form at a distance of 10 nm and 40 nm exceeds the critical voltage of the external electric field required to generate a transmembrane potential of 1V, which is sufficient for the electrical breakdown of bilayer lipid membranes.

Considering that under the conditions for which the electric field strength was calculated (pH 4.0), the cell surface is covered by many nanoparticles (as evidenced by the SEM image in the work of Schwegmann et al.^14^), each of which generates an electric field high voltage, it should be expected that destruction of cell membranes may occur.

After the destruction of the insulating barrier in the form of membranes, the electric field can act on the unprotected DNA molecule. The calculated strength of the electric field that a nanoparticle can generate inside the cell in the area where the DNA is located is sufficient to cause the collapse of the DNA. This means that under the exposure of TiO_2_ NPs in the absence of UV-A irradiation when photocatalysis does not occur and there is no ROS formation, cell death can occur due to electrical breakdown of bilayer lipid membranes and destruction of the DNA.

Photoactivation of TiO_2_ NPs by UV-A irradiation, as previously established, leads to protonation of the surface of nanoparticles.^35^ This means that with an increase in the duration of UV-A irradiation, the positive charge of their surface increases and, therefore, the electric field strength created by it as well. This gives reason to believe that when microorganisms are exposed to TiO_2_ NPs in combination with UV-A irradiation, cells undergo a stronger electric field of nanoparticles than without UV-A irradiation.

Protonation of the surface of TiO_2_ NPs during its exposure to UV-A irradiation can explain why, when cells are exposed to TiO_2_ NPs in combination with UV-A irradiation, cell inactivation occurs under pH conditions both in the pI region and above or below it (Figure 1), that is, when the surface of the nanoparticles is initially uncharged or charged negatively or positively,^13^ in contrast to the dark action of TiO_2_ NPs, for which the prerequisite is the presence of a positive charge initially.^11^

Apparently, when the surface of nanoparticles is protonated under the action of UV-A irradiation, then over time it is recharged to a positively charged one, if it was initially neutral or negatively charged. This assumption is confirmed by the presence of a lag period (a delay in the reduction of CFUs for some time) if experiments were carried out with such nanoparticles.^13^ Probably as soon as the value of the positive charge reaches some sufficient value after recharging, the CFUs begins to decrease. The recharging of the surface is indicated by previously obtained facts that, when cells of microorganisms are exposed to TiO_2_ NPs in combination with UVA irradiation, the pH value of the medium drops to values corresponding to an acidic medium, in which NPs are positively charged, although initially the medium was neutral^8^ or even alkaline,^13^ that is, when the TiO_2_ NPs were initially uncharged or negatively charged, respectively. The possible recharge of the surface under the action of UV-A irradiation was mentioned earlier.^13^

Although the facts indicate that when microorganisms are exposed to TiO_2_ NPs in combination with UV-A irradiation, the death of the latter is caused by both initially uncharged and positively and negatively charged nanoparticles, with a high degree of probability it can be argued that nanoparticles have a direct detrimental effect in cationic form.

The important role of the positive charge of TiO_2_ NPs in their antimicrobial action previously was pointed out in a number of works.^2,4,11,17,18,28^. At the same time, the role of the positive charge of TiO_2_ NPs was considered not as a source of an electric field, but as a necessary condition for ensuring close contact between the surfaces of cells and nanoparticles, which, according to the authors, is the optimal condition for the action of ROS, since they are formed precisely on the surface of nanoparticles during UV-A irradiation. Under conditions when the surface of TiO_2_ NPs is positively charged, such contact can be provided due to the electrostatic interaction between the cells and nanoparticles, since the surface of most bacterial cells in a wide pH range (from ~ 2.0 to ~ 11.0) is negatively charged.^11,14^ However, considering that TiO_2_ NPs can generate a high electric field the role of a positive charge as a source of this field seems to be more important. Moreover, if the inactivation of microorganisms does indeed occur as a result of the influence of an electric field, then it can be assumed that close contact between the surfaces of cells and nanoparticles, the importance of which was emphasized in a number of works, is not necessary, since the electric field can influence remotely. This conclusion is confirmed by the facts obtained by Kikuchi et al.^36^, where bacterial cells and a TiO_2_-based nanomaterial were separated by a semipermeable membrane that prevented them from contacting each other. It turned out that in this case, there was a decrease in the survival of bacterial cells also.

## 4. Conclusion

Thus, mathematical calculations show that cationic TiO_2_ NPs are capable of generating an electric field with a strength sufficient for the electrical breakdown of bacterial membranes. It seems very likely that the key mechanism responsible for cell death as a result of exposure to TiO_2_ NPs, both in combination with UV-A irradiation or without it, could be an electrical break-down of the energy-transforming CPM and destruction of DNA as a result of the electric field action, created by cationic TiO_2_ NPs.

It should be noted that energy-transforming membranes are present in gram-negative and gram-positive bacteria with a respiratory type of energy, in photosynthetic bacteria, as well as in mitochondria and chloroplasts of eukaryotes. This gives reason to believe that the electric field of cationic nanoparticles can have a destructive effect on the membranes of these objects.

It is likely that other cationic nanoparticles (AgNPs, ZnO NPs, MgO NPs, etc.) can also exert an antimicrobial effect through their electric field. The presence of a positive charge and sufficient electric field strength are important. To verify this assumption, the electric field strength of other cationic nanoparticles should be calculated.

## Supporting information

Supplementary Information contains calculation of the TiO2 na-noparticle electric field strength inside the E. coli K 12 cell at different distances f

## ASSOCIATED CONTENT

### Supplementary Information

Supplementary Information contains calculation of the TiO_2_ nanoparticle electric field strength inside the E. coli K 12 cell at different distances from the nanoparticle.

### AUTHOR INFORMATION

#### Present Addresses

Bach Institute of Biochemistry, Research Center of Biotechnology of the Russian Academy of Sciences. 33, bld. 2 Leninsky Ave., Moscow 119071, Russia.

### Author Contributions

The article as well as the Figures were made by the author.

### Notes

The author declare no competing financial interests.

## ACKNOWLEDGMENT

The author thanks the Bach Institute of Biochemistry, Research Center of Biotechnology of the Russian Academy of Sciences for the opportunity to do this work.

## REFERENCES

1. Huh, A. J.; Kwon, Y. J. “Nanoantibiotics”: A new Paradigm for Treating Infectious Diseases Using Nanomaterials in the Antibiotics Resistant Era. J Controll Release 2011, 156, 128–145. DOI:10.1016/j.jconrel.2011.07.002

2. Wang, L.; Hu, C.; Shao, L. The Antimicrobial Activity of Nanoparticles: Present Situation and Prospects for the Future. Int J Nanomedicine 2017, 12, 1227–1249. DOI:10.2147/IJN.S121956

3. Raghunath, A.; Perumal, E. Metal Oxide Nanoparticles as Antimicrobial Agents: a Promise for the Future. Int J Antimicrob Agents 2017, 49, 137–152. DOI:10.1016/j.ijantimicag.2016.11.011

4. Foster, H. A.; Ditta, I. B.; Varghese, S.; Steele, A. Photocatalytic Disinfection Using Titanium Dioxide: Spectrum and Mechanism of Antimicrobial Activity. Appl Microbiol Biotechnol 2011, 90, 1847–1868. DOI:10.1007/s00253-011-3213-7

5. Laxma Reddy, P. V.; Kavitha, B.; Kumar Reddy, P. A.; Kim, K. H. TiO_2_-based Photocatalytic Disinfection of Microbes in Aqueous Media: A Review. Environ Res 2017, 154, 296–303. DOI:10.1016/j.envres.2017.01.018

6. Uyguner Demirel, C. S.; Birben, N. C.; Bekbolet, M. A. Comprehensive Review on the Use of Second Generation TiO_2_ Photocatalysts: Microorganism Inactivation. Chemosphere 2018, 211, 420–448. DOI:10.1016/j.chemosphere.2018.07.121

7. Matsunaga, T.; Tomoda, R.; Nakajima, T.; Wake, H. Photoelectrochemical Sterilization of Microbial Cells by Semiconductor Powders. FEMS Microbiol Lett 1985, 29, 211–214. DOI:10.1111/j.1574-6968.1985.TB00864.X

8. Saito, T.; Iwase, T.; Horie, J.; Morioka, T. Mode of Photocatalytic Bactericidal Action of Powdered Semiconductor TiO_2_ on Mutans Streptococci. J Photochem Photobiol B 1992, 14, 369–379. DOI:10.1016/1011-1344(92)85115-b

9. Adams, L. K.; Lyon, D. Y.; Pedro, J. J.; Alvarez, P. J. J. Comparative Eco-Toxicity of Nanoscale TiO_2_, SiO_2_, and ZnO Water Suspensions. Water Res 2006, 40, 3527–3532. DOI:10.1016/j.watres.2006.08.004

10. Zhukova, L. V.; Kiwi, J.; Nikandrov, V. V. TiO_2_ Nanoparticles Suppress Escherichia coli Cell Division in the Absence of UV Irradiation in Acidic Conditions. Colloids Surf B 2012, 97, 240–247. DOI:10.1016/j.colsurfb.2012.03.010

11. Pagnout, C.; Jomini, S.; Dadhwal, M.; Caillet, C.; Thomas, F.; Bauda, P. Role of Electrostatic Interactions in the Toxicity of Titanium Dioxide Nanoparticles Toward Escherichia coli. Colloids Surf B 2012, 92, 315–321. DOI:10.1016/j.colsurfb.2011.12.012

12. Bae, H. S.; Lee, M. K.; Kim, W. W.; Rhee, C. K. Dispersion Properties of TiO_2_ Nano-Powder Synthesized by Homogeneous Precipitation Process at Low Temperatures. Colloids Surf A: Physicochem Eng Aspects 2003, 220, 169–177. DOI:10.1016/S0927-7757(03)00077-3

13. Gumy, D.; Morais, C.; Bowen, P.; Pulgarin, C.; Giraldo, S.; Hajdu, R.; Kiwi. J. Catalytic Activity of Commercial of TiO_2_ Powders for the Abatement of the Bacteria (*E. coli*) under Solar Simulated Light: Influence of the Isoelectric Point. Appl Catal, B. 2006, 63, 76–84. DOI:10.1016/j.apcatb.2005.09.013

14. Schwegmann, H.; Ruppert, J.; Frimmel, F.H. Influence of the pH-Value on the Photocatalytic Disinfection of Bacteria with TiO_2_ – Explanation by DLVO and XDLVO Theory. Water Res 2013, 47, 1503–1511. DOI:10.1016/j.watres.2012.11.030

15. Sunada, K.; Watanabe, T.; Hashimoto, K. Studies on photokilling of bacteria on TiO_2_ thin film. J Photochem Photobiol A 2003, 156, 227–233. DOI:10.1016/s1010-6030(02)00434-3

16. Ganguly, P.; Byrne, C.; Breen, A.; Pillai, S.C. Antimicrobial Activity of Photocatalysts: Fundamentals, Mechanisms, Kinetics and Recent Advances. Appl Catal B 2018, 225, 51–75. DOI:10.1016/j.apcatb.2017.11.018

17. Mileyeva-Biebesheimer, O. N.; Zaky, A.; Gruden, C. L. Assessing the Impact of Titanium Dioxide and Zinc Oxide Nano-particles on Bacteria Using a Fluorescent-Based Cell Membrane Integrity Assay. Environ Eng Sci 2010, 27, 329–335. DOI:10.1089/ees.2009.0332

18. Sohm, B.; Immel, F.; Bauda, P.; Pagnout, C. Insight into the Primary Mode of Action of TiO_2_ Nanoparticles on Escherichia coli in the Dark. Proteomics 2015, 15, 98–113. DOI:10.1002/pmic.201400101

19. Sale, A. J. H.; Hamilton, W. A. Effects of High Electric Fields on Microorganisms. I. Killing of Bacteria and Yeasts. Biochim. Biophys. Acta 1967, 148, 781–788. DOI:10.1016/0304-4165(67)90052-9

20. Hülsheger, H.; Potel, J.; Niemann, E.-G. Electric Field Effects on Bacteria and Yeast Cells. Radiat Environ Biophys 1983, 22, 149–162. DOI:10.1007/BF01338893

21. Kinosita, Jr. K.; Tsong, T. Y. Formation and Resealing of Pores of Controlled Size in Human Erythrocyte Membrane. Nature 1977, 268, 438–441. DOI: 10.1038/268438a0

22. Sale, A. J. H.; Hamilton, W. A. Effects of high electric fields on microorganisms. III. Lysis of erythrocytes and protoplasts. Biochim. Biophys. Acta 1968, 43, 37–43. DOI:10.1016/0005-2736(68)90030-8

23. Weaver, J. C.; Chizmadzhev, Y. Theory of Electroporation: A Review. Bioelectoch Bioener 1996, 41, 135–160. DOI:10.1016/S0302-4598(96)05062-3

24. Chang, D. C.; Reese, T. S. Changes in Membrane Structure Induced by Electroporation as Revealed by Rapid-Freezing Electron Microscopy. Biophys J 1990, 58, 01–12. DOI:10.1016/S0006-3495(90)82348-1

25. Vladimirov, Yu. A.; Biological Membranes and Non-Programmed Cell Death. Soros Education journal 2000, 6, 2–9.

26. Schlegel, H. G. Allgemeine Mikrobiologie; 6th ed, Schmidt, K., Ed; Georg Thieme Verlag: Stuttgart New York, 1985, 566 p. DOI:10.1002/star.19810331114

27. Skulachev, V. P., Bogachev A. V., Kasparinsky F. O. Membrane bioenergetics: Pevak, E. A., Ed.; Moscow University Press: Moscow, 2010. 368 p.

28. Gogniat, G.; Thyssen, M.; Denis, M.; Pulgarin, C.; Dukan, S. The Bactericidal Effect of TiO_2_ Photocatalysis Involves Adsorption onto Catalyst and the Loss of Membrane Integrity. FEMS Microbiol Lett 2006, 258, 18–24. DOI:10.1111/j.1574-6968.2006.00190.x

29. Tang, J.; Du, N.; Doyle, P.S. Compression and Self-Entanglement of Single DNA molecules under Uniform Electric Field. PNAS 2011, 108, (39), 16153–16158. DOI:10.1073/pnas.1105547108

30. Nomura, N.; Abe, S.; Uchida, I.; Abe, K.; Koga, H.; Katsuki, S.; Namihira, T.; Akiyama, H.; Takano, H.; Abe, S.I. Direct Deformation of DNA Using Intense Burst RF Electric Field. 27^th^ International Power Modulator Symposium/2006 High Voltage Workshops 2006. Proceedings of the 27^th^ International Power Modulator Symposium and 2006 High Voltage Workshop, pp. 490–493. DOI:10.1109/MODSYM.2006.365291

31. Zhou, C.; Riehn, R. Collapse of DNA under Alternating Electric Fields. Phys Rev E Stat Nonlin Soft Matter Phys 2015, 92, 012714 DOI:10.1103/PhysRevE.92.012714

32. Holmberg, J. P.; Ahlberg, E.; Bergenholtz, J.; Hassellov, M.; Abbas, Z. Surface Charge and Interfacial Potential of Titanium Dioxide Nanoparticles: Experimental and Theoretical Investigations. J Colloid Interface Sci 2013, 407, 168–176. DOI:10.1016/j.jcis.2013.06.015

33. Silhavy, T. J.; Kahne, D.; Walker, S. The Bacterial Cell Envelope. Cold Spring Harb Perspect Biol 2010, 2: a000414. DOI:10.1101/cshperspect.a000414

34. Matias, V. R. F.; Al-Amoudi, A.; Dubochet, J.; Beveridge, T. J. Cryo-Transmission Electron Microscopy of Frozen-Hydrated Sections of Escherichia coli and Pseudomonas aeruginosa. J Bacteriol 2003, 185, (20), 6112–6118. DOI: 10.1128/JB.185.20.6112

35. Wang, Z-S.; Yamaguchi, T.; Sugihara, H.; Arakawa, H. Significant Efficiency Improvement of the Black Dye-Sensitized Solar Cell through Protonation of TiO_2_ Films. Langmuir 2005, 21, 4272–4276. DOI:10.1021/la050134w

36. Kikuchi, Y.; Sunada, K.; Iyoda, T.; Hashimoto, K.; Fujishima, A. Photocatalytic Bactericidal Effect of TiO_2_ Thin Films: Dynamic View of the Active Oxygen Species Responsible for the Effect. J Photochem Photobiol A, 1997, 106, 1–3, 51–56. DOI:10.1016/s1010-6030(97)00038-5

