## Supplementary Information contains calculation of the TiO2 na-noparticle electric field strength inside the E. coli K 12 cell at different distances f for "The electric field generated by cationic TiO_2_ nanoparticles can be their key antimicrobial factor": Supplementary Information.pdf

Lyudmila V. Zhukova*

Bach Institute of Biochemistry, Research Center of Biotechnology of the Russian Academy of Sciences. 33, bld. 2 Leninsky Ave., Moscow 119071, Russia

Here are the calculations of the $TiO_2$ nanoparticle surface area, surface charge and electric field strength inside the E. coli K 12 cell at different distances from the nanoparticle.

$S = 4\pi R^2$

Where S - the surface area of the nanoparticle; R - the radius of the nanoparticle (40 nm); π - a mathematical constant equal to 3.14.

$S = 4 \times 3{,}14 \times (40 \times 10^{-9}) = 20096 \times 10^{-18} = 2{,}0096 \times 10^{-14}$ ($m^2$)

$Q = S \times \sigma$

where Q - the charge of the nanoparticle; S - the surface area of the nanoparticle ($2{,}0096 \times 10^{-14}$ ($m^2$)); σ - the surface charge density (0.1 $C/m^2$).

$Q = 2{,}0096 \times 10^{-14} \times 0.1 = 2{,}0096 \times 10^{-15}$ (C)

$$E = \frac{1}{4\pi\varepsilon_0} \times \frac{Q}{\varepsilon \times (R+L)^2}$$

Where E - electric field strength

Q - the charge of the nanoparticle ($2{,}0096 \times 10^{-15}$ C);

R - the radius of the nanoparticle ($40 \times 10^{-9}$ m);

L - the distance at which the electric field strength is determined ($10 \times 10^{-9}$ m; $40 \times 10^{-9}$ m; $460 \times 10^{-9}$ m)

$\varepsilon_o$ - electric constant equal to $8.85 \times 10^{-12}$ ($CN^{-1}$ $m^{-2}$);

ε - dielectric constant of the medium equal to 80 (dimensionless value).

π - a mathematical constant equal to 3.14.

1. $L = 10 \times 10^{-9}$ m

$$E = 9 \times 10^9 \times \frac{2.0096 \times 10^{-15}}{80 \times [(40+10) \times 10^{-9}]^2} = 9.0432 \times 10^7 \text{ (N/C)} = (\sim 10^8) \text{ (V/m)}$$

2. $L = 40 \times 10^{-9}$ m

$$E = 9 \times 10^9 \times \frac{2.0096 \times 10^{-15}}{80 \times [[(40+40) \times 10^{-9}]]^2} = 3.5325 \times 10^7 \text{ (N/C)} = (\sim 10^7) \text{ (V/m)}$$

3. $L = 460 \times 10^{-9}$ m

$$E = 9 \times 10^9 \times \frac{2.0096 \times 10^{-15}}{80 \times [(40+460) \times 10^{-9}]^2} = 9.0432 \times 10^5 \text{ (N/C)} = (\sim 10^6) \text{ (V/m)}$$
